# Improving Molecular Docking Pose Recovery through Targeted Side-Chain Monte Carlo Sampling of an Apo Structure

**DOI:** 10.64898/2026.10.28.762659

**Authors:** Pavan Subramani, Mrinal Shekhar

**Affiliations:** Research Science Institute, Cambridge, MA, USA; Center for Development of Therapeutics, Broad Institute of MIT and Harvard, Cambridge, MA, USA

**Keywords:** Molecular docking, molecular dynamics, improved sampling, accurate docking

## Abstract

Receptor flexibility complicates molecular docking when only an unbound (apo) protein structure is available. Local side-chain rearrangements can alter binding-pocket accessibility and ligand complementarity, motivating approaches that sample these motions efficiently. Here, we evaluate a targeted Markov Chain Monte Carlo (MCMC) protocol that combines local and larger side-chain torsional proposals with periodic molecular dynamics (MD) relaxation under harmonic backbone restraints. Receptor ensembles generated from an apo structure were evaluated against 859 experimental complexes from the OpenBind 2A protease benchmark and compared with the apo receptor and ensembles generated by restrained and unrestrained MD. With regular-space representative selection, MCMC recovered a ligand pose within 2 Å of the experimental pose for 74.4% of complexes, compared with 51.0% for apo and 59.5% for restrained MD. Median minimum ligand heavy-atom RMSD decreased from 1.96 Å for apo and 1.82 Å for restrained MD to 1.62 Å for MCMC. The largest gains occurred for complexes with greater side-chain displacement relative to pocket backbone displacement, consistent with the motions targeted by the protocol. MCMC representatives retained pocket geometries within the experimentally observed range, whereas unrestrained-MD representatives exhibited reduced pocket free volume and poor recovery. These findings support targeted side-chain sampling as a practical approach to generating apo-derived docking ensembles. The reported improvements measure best-case pose coverage across generated poses and receptors; their extension to prospective pose ranking and virtual screening remains to be established.

## Introduction

Structure-based virtual screening is widely used in modern computer-aided drug design (CADD). Its success depends on accurately predicting binding poses and scoring and ranking small-molecule candidates docked from large chemical libraries. However, docking into unbound (apo) crystal or cryo-EM structures without accounting for receptor flexibility can yield inaccurate poses, as side-chain rotamers that occlude the binding pocket may clash with potential ligands. Receptor conformations that accommodate ligand binding are therefore essential for effective screening, and ensembles of such conformations can help account for binding-site flexibility [1].

These challenges motivate development of algorithms to elucidate receptor conformations that are amenable to docking. In recent times, structure prediction and ensemble generating approaches, including Alphafold [2] and BioEmu [3], offer alternate approaches towards generating these receptor conformations. Their utility for virtual screening, however, depends on whether the resulting structures capture binding-pocket geometries compatible with ligand binding. Generating such conformations remains a particular challenge when the relevant pocket rearrangements are unknown or poorly characterized [4]. Moreover, these AI tools underperform when there is no background knowledge about the target protein or what movements occur to allow for ligand binding.

Alternatively, molecular dynamics (MD) serves as a physics-based method for sampling receptor conformations; however, spontaneous transitions between apo and holo geometries remain infrequent on computationally accessible timescales.

Enhanced sampling approaches address this limitation through different strategies. Metadynamics applies a history-dependent bias along selected collective variables [5]. Replica exchange facilitates exploration through exchanges between simulations conducted under different conditions [6]. Weighted ensemble methods redistribute sampling effort among trajectory segments. The effectiveness and computational cost of these approaches depend on the system, sampling parameters, and observables of interest [7]. As an alternative to such extensive, ligand-independent sampling, induced-fit docking instead adapts receptor geometry directly around individual candidate ligands, generating conformations specific to those docking calculations. However, because the computational cost of dynamically optimizing the receptor must be incurred for every individual compound, induced-fit docking remains intractable for large-scale virtual screening [8].

Overcoming these limitations requires a method that bypasses global protein sampling to focus exclusively on local binding-site rearrangements. Monte Carlo (MC) sampling offers a highly direct method for proposing these targeted transitions between side-chain conformations. Previous research successfully combined diffusion-based proposals with Markov Chain Monte Carlo sampling of alanine dipeptide [9], utilizing Metropolis-Hastings acceptance to target the Boltzmann distribution [10][11].

For receptor generation, the distinct objective is to produce a compact, diverse ensemble of low-energy pocket conformations suitable for docking. This approach prioritizes exploring physically valid binding-site geometries over determining their exact thermodynamic probabilities, provided the underlying sampling rules remain rigorous.

To investigate targeted side-chain sampling, we focused on the EV-A71 2A protease, a protein implicated in enterovirus infection and hand, foot, and mouth disease [12]. The OpenBind dataset provided experimentally determined receptor–ligand complexes alongside an apo reference for this system [13]. These robust structural benchmarks enable the systematic evaluation of docking pose recovery across numerous ligands and allow us to analyze how experimental pocket rearrangements impact the overall benefit of receptor sampling. The availability of a large structural benchmark for this target is further described by the OpenBind consortium [14].

Here, we introduce a Monte Carlo protocol that combines local torsional perturbations with periodic MD relaxation under backbone restraints. We compare ensembles generated by this protocol with the apo structure and with restrained and unrestrained MD ensembles. Two clustering strategies test whether selecting frequently sampled conformations or emphasizing conformational diversity is more useful for docking. We then relate docking performance to experimental backbone and side-chain displacement and to pocket geometry. The study evaluates the utility of targeted side-chain sampling for pose recovery on one protein target; it does not establish prospective screening enrichment or equilibrium sampling accuracy.

## Method

### Benchmark dataset and structure preparation

Experimental structures of EV-A71 2A protease were obtained from the OpenBind dataset [14]. The structural analysis included 925 prepared holo structures, and docking performance was evaluated for 859 dockable complexes. Of these complexes, 858 had a defined backbone-to-side-chain displacement ratio and were included in the stratified analysis. Docking complexes were selected using the OpenBind metadata by requiring a valid prepared structure and excluding suspected artifacts. Covalent complexes were also excluded when the corresponding metadata annotation was available. Experimental ligand coordinates served as the reference poses for docking evaluation. Binding-affinity measurements were not used as the primary endpoint.

### Binding-pocket definition

A fixed set of pocket residues was used throughout receptor sampling, trajectory clustering, and pocket structural analyses. The pocket was defined from an experimental receptor–ligand complex by selecting every protein residue containing at least one atom within 5.0 Å of a reference ligand atom. The resulting 21 residues comprised of resid 21, 39, 84, 85, 87, 89, 90, 93, 95, 105, 106, 107, 109, 110, 124, 125, 126, 127, 128, 129, and 132. This residue set was retained across complexes rather than redefined for each ligand. The experimental ligand was used to identify the pocket but was absent during MCMC and MD receptor sampling.

### Molecular Model and Simulation Settings

Structures were prepared using PDBFixer [15] to identify and model missing residues and atoms, strip heterogens and solvent, and add hydrogens appropriate for pH 7.0. All simulations were performed in OpenMM [16] using the Amber ff14SB force field [17] and the GBn2 implicit solvent model [18] at 300 K. Dynamics were propagated using a Langevin middle integrator [19] with a friction coefficient of 1.0 ps^−1^ and a 2 fs timestep.

While the MCMC and MD protocols shared the same force field parameters, solvent model, temperature, friction coefficient, and timestep, their OpenMM system configurations differed slightly. The MCMC system employed untruncated nonbonded interactions (NoCutoff), omitted bond constraints, and disabled center-of-mass motion removal. In contrast, the MD control simulations utilized a 2.0-nm nonbonded cut-off (CutoffNonPeriodic), constrained all bonds involving hydrogen, set hydrogen masses to 1.5 amu, and enabled center-of-mass motion removal. Additionally, the MD simulations were explicitly executed using CUDA mixed precision.

### Markov Chain Monte Carlo

The technique we created was Markov Chain Monte Carlo for protein side chain sampling. This was chosen because it allows for extremely quick sampling because we can propose lots of moves cheaply. For each proposal, one eligible side-chain *χ* torsion from the fixed pocket was selected uniformly at random. Selection was uniform over eligible torsions rather than residues, so residues containing multiple eligible *χ* torsions could be selected through more than one torsional degree of freedom. With 70% probability, a local rotation was drawn uniformly from −30° to +30°. With 30% probability, a large rotamer-jump proposal was made by selecting the positive or negative direction with equal probability and drawing the magnitude as (120° + *U* (−20°, 20°)), corresponding to a magnitude between 100° and 140°.

This method was chosen to allow for both small- and large-scale exploration. We then apply the move by rotating the atom set around the bond and evaluate the energy of the entire system in the new state. The potential energy of the complete OpenMM system, including the backbone restraint, was evaluated before and after the proposed torsional rotation.

A proposed move with Δ*U* = *U*_new_ − *U*_old_ was accepted with probability *P*_acc_ = min[1, exp(−*β*Δ*U*)], where *β* = (*k*_B_*T*)^−1^. “Downhill” moves to lower-energy states were therefore always accepted, whereas uphill moves were accepted probabilistically according to their energy increase. Rejected proposals were reverted to the previously accepted coordinates. This is the Metropolis filter [10]. It was used to promote the sampling of physically plausible simulation states.

Every 2000 accepted moves, we run 1 picosecond of classical MD to allow the backbone and side chains to relax around the new state. Immediately before each relaxation, velocities were newly initialized from a 300-K distribution. Velocities were therefore not propagated continuously between separate MD relaxation bursts.

This was all done with a protein backbone restraint. A harmonic spring (k = 1000 kJ/mol/nm^2^) tethers every backbone heavy atom (N, CA, C, O) to its position in the prepared starting receptor. The backbone was harmonically restrained rather than fixed, allowing limited backbone relaxation while strongly favoring the starting apo geometry. An illustration of how this method worked is shown in fig. 1.

**Figure 1:**
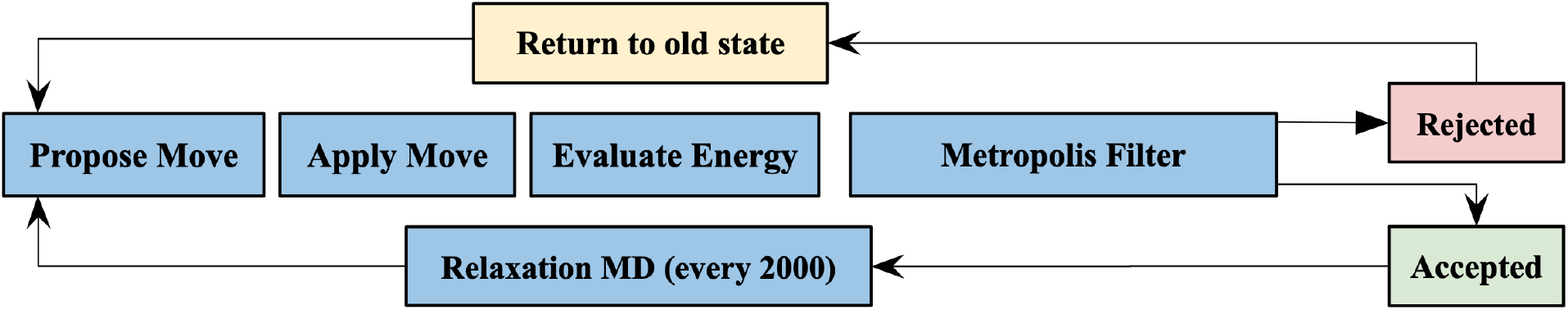
MCMC Sampling Loop. We propose the move, apply it by rotating the atoms, evaluate the energy at the new state, and pass it through a Metropolis criterion to promote physically plausible states.

We generated a trajectory of the MCMC method with random seed 0 and 50,000,000 attempted Monte Carlo proposals. We saved coordinates every 1000 attempted proposals, which created 50,000 stored frames. Trajectory output was indexed by attempted proposal number rather than accepted-move number. Because of this, if the proposal at a saved step was rejected, the current accepted configuration was written and consecutive saved frames could be identical.

### Controls

We initiated two ligand-free implicit-solvent MD controls from the same apo starting structure: a 1 microsecond unrestrained trajectory and a 200 ns backbone-restrained trajectory, both starting from the apo structure. For each newly initialized MD simulation, the structure was energy-minimized for up to 5000 iterations, velocities were initialized at 300 K, and the system was equilibrated for 0.5 ns before production. The restrained MD control used the same 1000 kJ mol^−1^ nm^−2^ harmonic positional restraint on backbone N, C*α*, C, and O atoms used by the MCMC sampler. Production coordinates were saved every 12.5 ps. The 1 microsecond unrestrained trajectory therefore contained 80,000 saved production frames and the 200-ns restrained trajectory contained 16,000 frames.

We compared docking to the representative structures generated by the MCMC technique described above to docking to the singular apo structure, docking to structures from 1 microsecond of MD, and docking to structures that came from the backbone-restrained MD.

### Analysis

#### Trajectory clustering and receptor selection

Trajectory frames were represented using the *χ*_1_ side-chain dihedral angles of pocket residues for which *χ*_1_ was defined. Each angle was encoded as [sin(*χ*_1_), cos(*χ*_1_)] to account for angular periodicity. The resulting feature dimensions were treated equally, and ordinary Euclidean distance in the sin/cos feature space was used for clustering.

Two different clustering algorithms were compared. We first clustered by population-based clustering, which used k-means with k=8, and retained one observed trajectory frame nearest each cluster centroid. The second algorithm was regular space clustering, which was chosen because it selects to create conformational diversity regardless of population.

Regular-space clustering greedily selected conformations separated by a specified minimum Euclidean distance in *χ*1 sin/cos feature space. The number of representatives therefore emerged from the distance criterion rather than being specified directly. The frame order used for regular-space selection was randomized using seed 0.

For population-based clustering, the observed trajectory frame nearest the k-means centroid was selected as the representative receptor. For regular-space clustering, each retained regular-space center was itself an observed trajectory frame and was used as the representative receptor.

#### Molecular docking and pose evaluation

To evaluate the method, we docked ligands into the representative receptor structures using SMINA (a fork of AutoDock Vina) [20]. Before docking, each sampled receptor was superposed onto the experimental receptor for the ligand being evaluated using matched pocket C*α* atoms, placing all receptor conformations in the corresponding crystal coordinate frame. For each experimental complex, the docking search box was constructed from that complex’s experimental reference ligand coordinates. The box center was the arithmetic mean of the ligand atom coordinates, and each dimension was the ligand extent plus 6 Å of padding on each side, with a minimum edge length of 16 Å. We docked with an exhaustiveness of 8 and generated up to 9 poses per run, all with a fixed random seed of 0.

Docking accuracy was quantified using the minimum symmetry-aware heavy-atom RMSD to the experimental ligand pose among all generated docking poses. For receptor ensembles, this minimum was further taken over all receptor conformations in the receptor set. Thus, the reported metric measures best-case pose-recovery coverage. A pose was counted as recovered if it was within a 2 Å threshold. The results are also reported within 3 Å and with a median minimum RMSD. Individual docking or receptor-alignment failures were skipped. If no receptor in a receptor set produced a valid result, that method’s value for the complex was recorded as missing.

To evaluate how much of the result was due to the accuracy of the docking method (SMINA), we also self-docked the ligands back into the experimental poses to see how often they were recovered. The gap between this self-docking recovery rate and 100% is due to docking error.

#### Experimental apo-to-holo structural analysis

To characterize the structural regime in which MCMC improves docking, experimental holo structures were compared with the apo reference. Each holo structure was globally aligned to the apo structure using common protein C*α* atoms. At least 20 common protein C*α* atoms were required. Pocket backbone displacement was quantified as the RMSD of pocket C*α* atoms after this alignment. At least eight common pocket C*α* atoms were required. Side-chain displacement was quantified as the heavy-atom RMSD of pocket side chains after local alignment on the common pocket backbone atoms (N, C*α*, C, and O), thereby separating side-chain rearrangement from backbone displacement. At least 20 common pocket-backbone atoms and 12 common pocket side-chain atoms were required for the corresponding calculations. Symmetry-equivalent terminal atoms were accounted for when calculating side-chain RMSD. For explicitly handled symmetry-equivalent side-chain atom assignments, both the identity assignment and permitted symmetry-swapped assignments were evaluated, and the assignment producing the smallest residue-level squared deviation was retained. A backbone-to-side-chain motion ratio was then defined as pocket C*α* RMSD divided by pocket side-chain RMSD. Lower ratios correspond to transitions dominated by side-chain rearrangement. If the side-chain RMSD was effectively zero, the ratio was treated as undefined.

#### Structural-regime and statistical analysis

Docking recovery was stratified into six equal-count bins of this ratio, and recovery was defined using the same ligand pose RMSD ≤ 2Å criterion used for the primary docking analysis. The 858 complexes with a defined ratio were therefore divided into six bins of 143 complexes each. For each method and ratio bin, the recovery proportion was calculated, and 95% Wilson score confidence intervals were used to quantify uncertainty in the bin-wise recovery proportions [21]. Separately, to compare MCMC directly with each comparator within each bin, the paired difference in recovery probability was estimated using 10,000 paired bootstrap resamples with random seed 0 [22]. Complex indices were sampled with replacement and applied jointly to both methods so that pairing by experimental complex was preserved. The 95% confidence interval for the recovery difference was defined by the 2.5th and 97.5th percentiles of the paired-bootstrap distribution.

We also applied the Wilcoxon signed-rank test to compare between methods by pairing per ligand [23]. Overall paired RMSD comparisons were performed using scipy.stats.wilcoxon on per-complex RMSD values [24]. The analysis did not implement a multiple-comparison correction.

#### Pocket SASA and free-volume analysis

To assess whether the conformations selected by clustering reproduced experimental pocket geometry, pocket solvent-accessible surface area (SASA) and pocket free volume were calculated for the regular-space and population-based representatives from MCMC, restrained MD, and unrestrained MD, as well as the apo structure and experimental holo structures. Pocket SASA was calculated for the fixed pocket residue set. SASA was calculated with MDTraj’s [25] Shrake-Rupley implementation in atom mode using a 1.4 Å solvent probe and 960 sphere points per atom. Hydrogen atoms were excluded, and the SASA values of the heavy atoms belonging to the fixed 21-residue pocket were summed.

Pocket free volume was calculated after global C*α* alignment to the experimental A71EV2A-x5584b holo reference. The 24 heavy atoms of its reference ligand defined a fixed inclusion region consisting of the union of 4.0Å radius spheres centered on the ligand atoms.

The region was discretized using a 0.5Å Cartesian grid. Grid points lying within the van der Waals radius of any aligned protein heavy atom were classified as occupied, and free volume was calculated from the number of remaining grid points. The ligand defined the spatial region but was not included as an occupying object; no flood-fill or solvent-connectivity criterion was applied, so this quantity represents total unoccupied volume within the fixed ligand-centered region rather than necessarily a single connected cavity.

## Results and Discussion

### Experimental Pocket Heterogeneity and Improved Pose Recovery with MCMC

The selected system exhibited substantial conformational variation between apo and holo binding pockets. Following the filtering procedure described in the Methods, 858 complexes had a defined backbone-to-side-chain displacement ratio and were retained for subsequent analysis. Superposition of the experimental structures revealed heterogeneous pocket conformations (fig. 2A), with contributions from both backbone and side-chain rearrangements relative to the apo reference (fig. 2B). Most structures fell below the line of equal backbone and side-chain RMSD, indicating that side-chain displacement generally exceeded pocket C*α* displacement under the respective alignment conventions.

**Figure 2:**
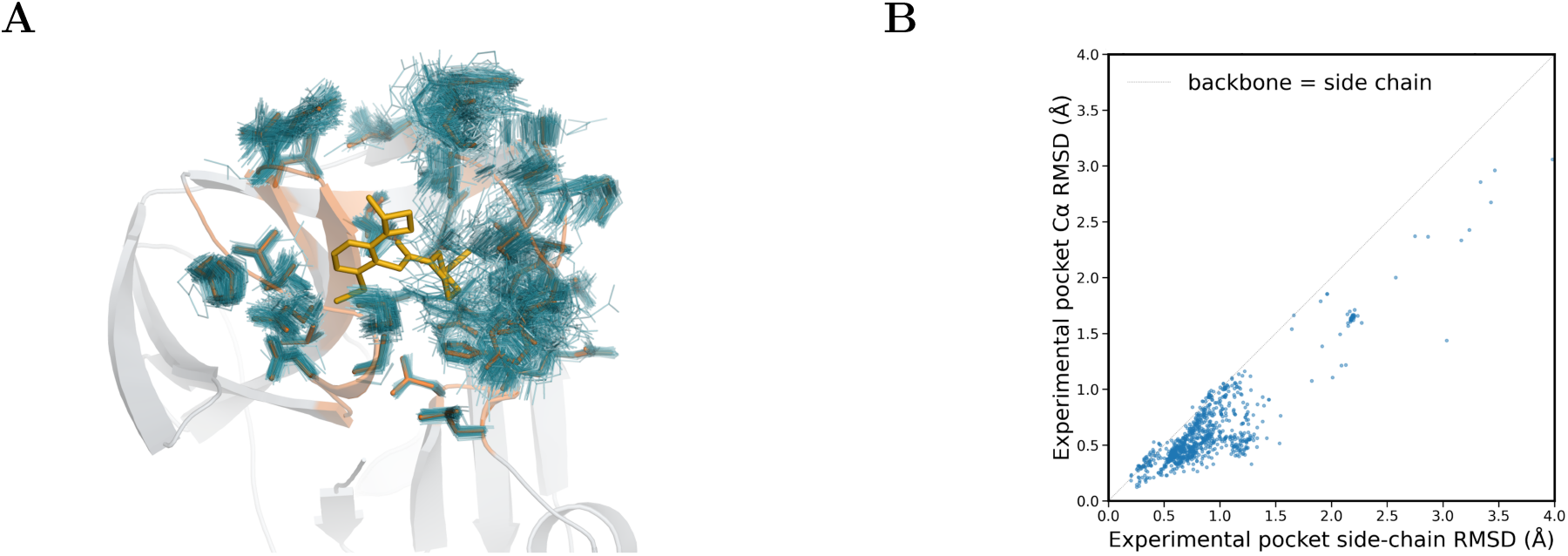
**(A)** Superposition of the experimental structures, where each stick is a pocket side chain from the experimental structure. **(B)** Experimental pocket C*α* RMSD and side-chain heavy-atom RMSD relative to the apo reference.

To assess whether this pocket heterogeneity could be addressed through receptor sampling, we compared docking into the apo structure with docking into computationally generated ensembles. Representative receptors were selected using regular-space clustering, which emphasizes diversity in pocket side-chain conformations. Among the apo-derived receptor sets, the resulting Monte Carlo ensemble achieved the highest pose recovery (table 1). At the 2 Å threshold, recovery increased from 51.0% for apo to 74.4% for Monte Carlo, or a relative increase of 45.8%. The regular-space restrained-MD ensemble achieved 59.5% recovery, giving the Monte Carlo method a relative 25.0% increase when compared to it. Median minimum RMSD decreased from 1.96 Å for apo and 1.82 Å for restrained MD to 1.62 Å for Monte Carlo.

**Table 1:**
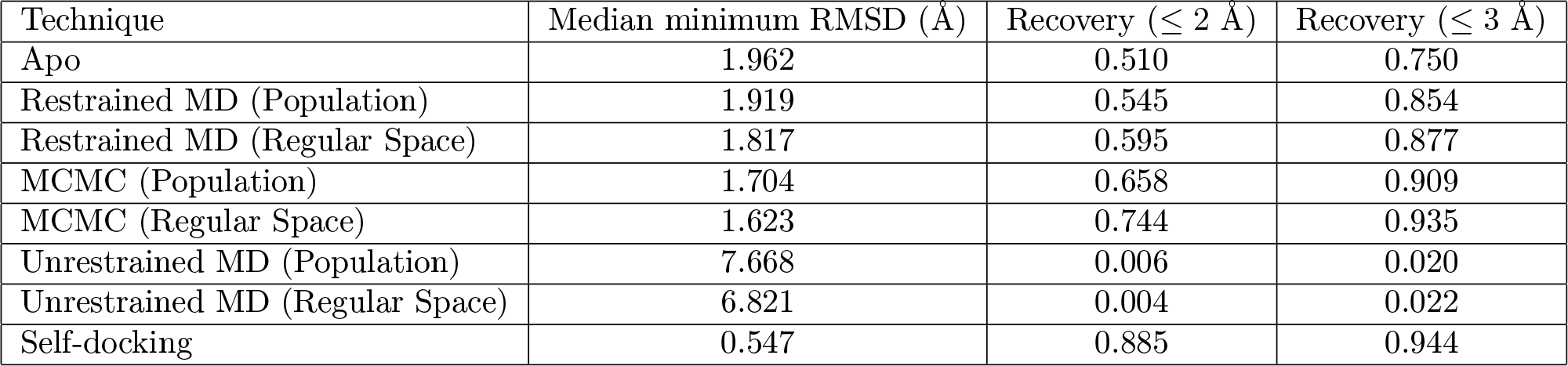
Best-case docking pose recovery across receptor-generation and representative-selection strategies. For each method–complex pair, the minimum symmetry-aware ligand heavy-atom RMSD was taken over all generated poses and, where applicable, all receptor conformations. Recovery is reported as the fraction of complexes with a minimum RMSD at or below the specified threshold.

| Technique | Median minimum RMSD (Å) | Recovery (≤ 2 Å) | Recovery (≤ 3 Å) |
|---|---|---|---|
| Apo | 1.962 | 0.510 | 0.750 |
| Restrained MD (Population) | 1.919 | 0.545 | 0.854 |
| Restrained MD (Regular Space) | 1.817 | 0.595 | 0.877 |
| MCMC (Population) | 1.704 | 0.658 | 0.909 |
| MCMC (Regular Space) | 1.623 | 0.744 | 0.935 |
| Unrestrained MD (Population) | 7.668 | 0.006 | 0.020 |
| Unrestrained MD (Regular Space) | 6.821 | 0.004 | 0.022 |
| Self-docking | 0.547 | 0.885 | 0.944 |

The advantage persisted at the 3 Å threshold, where recovery was 93.5% for Monte Carlo, 87.7% for restrained MD, and 75.0% for apo. The cumulative recovery curves showed that the comparison was not confined to the primary 2Å threshold (fig. 3B).

**Figure 3:**
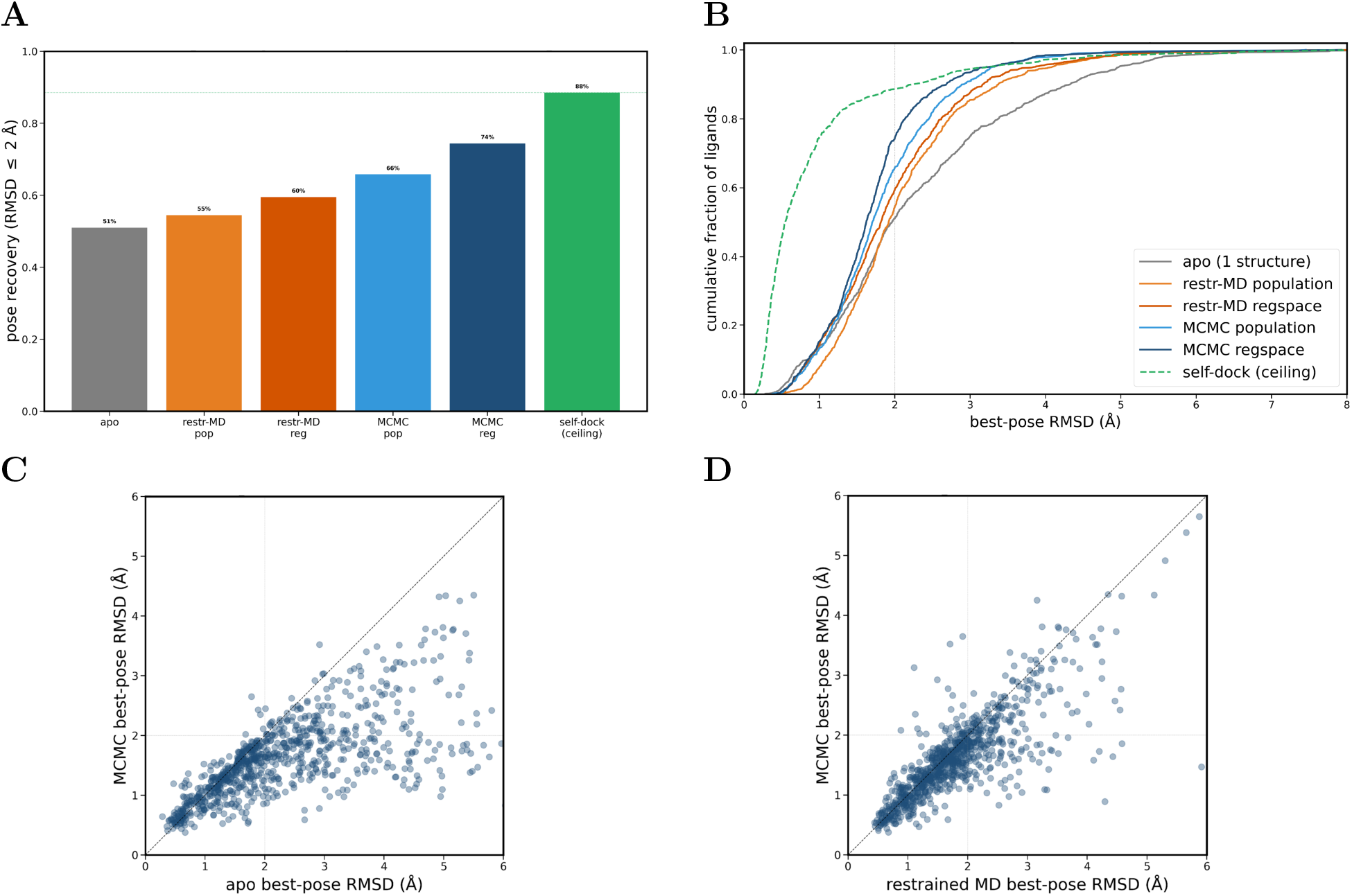
Comparison of docking performance across methods. **(A)** Percent recovery of each docking technique relative to the self-docking reference. **(B)** Fraction of ligands recovered as a function of the RMSD threshold, showing how docking success varies with the definition of a successfully recovered pose. **(C)** Comparison of best-pose RMSD between MCMC and apo structure. **(D)** Comparison of best-pose RMSD between MCMC and restrained MD.

Monte Carlo yielded a lower minimum RMSD than apo for 675 of 859 complexes and a lower minimum RMSD than restrained MD for 580 of 859 complexes (fig. 3C,D). Paired Wilcoxon tests supported differences in the corresponding RMSD distributions, with p < 10^−89^ for the comparison to apo and p < 10^−34^ for the comparison to restrained MD.

Self-docking recovered 88.5% of complexes within 2 Å and yielded a median minimum RMSD of 0.55 Å. The regular-space Monte Carlo and restrained-MD ensembles therefore achieved 84.1% and 67.2% of the observed self-docking recovery rate, respectively.

The effects of representative-selection strategy and ensemble size are examined in the next section.

### Representative Selection Influences Ensemble Performance

Population-based and regular-space selection emphasize different regions of the sampled conformational space. Population-based selection retains representatives from the most populated clusters, whereas regular-space selection can retain configurations farther from population centers. This distinction affected docking performance for both MCMC and restrained MD.

For MCMC, regular-space selection increased recovery within 2 Å from 65.8% to 74.4% and decreased median minimum RMSD from 1.70 to 1.62 Å when compared to population-based selection (table 1). For restrained MD, recovery increased from 54.5% to 59.5%, while median minimum RMSD decreased from 1.92 to 1.82 Å. Recovery within 3 Å also improved, increasing from 90.9% to 93.5% for MCMC and from 85.4% to 87.7% for restrained MD.

These comparisons involved different numbers of representative receptors. Population-based selection retained eight representatives for each method, whereas regular-space selection retained 15 for MCMC, seven for restrained MD, and 14 for unrestrained MD. The MCMC improvement therefore reflects changes in both representative selection and ensemble size. Restrained MD performed better with regular-space selection than with population-based selection despite using one fewer representative, indicating that receptor selection can affect recovery independently of an increase in receptor count.

MCMC also outperformed restrained MD when both methods used eight population-based representatives: recovery within 2 Å was 65.8% and 54.5%, respectively. Thus, the MCMC advantage was present under both selection strategies, although its magnitude depended on the resulting receptor ensemble.

### Recovery Gains Depend on the Character of Pocket Rearrangement A B

Having established the effect of representative selection, we examined whether the experimental pattern of pocket rearrangement was associated with the benefit of MCMC. The 858 complexes with a defined pocket C*α*-to-side-chain RMSD ratio were divided into six equal-count bins of 143 complexes. Lower ratios indicate greater side-chain displacement relative to pocket backbone displacement under the specified alignment conventions.

The regular-space MCMC ensemble achieved higher recovery than apo and restrained MD in all six bins (fig. 4A). Its largest advantage over restrained MD occurred at ratios below approximately 0.70, where recovery gains were approximately 19–20 percentage points. The advantage generally decreased as backbone displacement became larger relative to side-chain displacement, although the relationship was not strictly monotonic.

**Figure 4:**
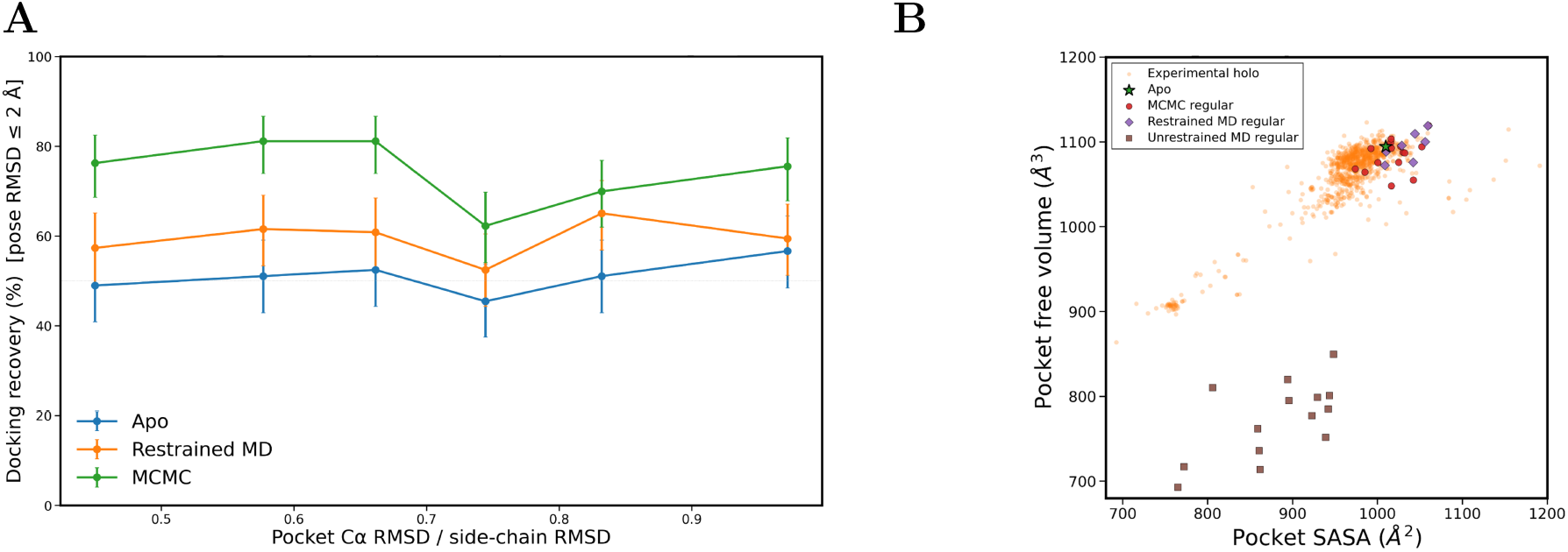
**(A)** Docking recovery rate as a function of the ratio of pocket C*α* RMSD to side-chain RMSD for the apo structure, restrained MD regular-space ensemble, and MCMC regular-space ensemble. Points represent six equal-count bins; error bars show 95% Wilson score confidence intervals for the bin-wise recovery proportions. Recovery was defined as ligand pose RMSD ≤ 2 Å. **(B)** Pocket SASA and pocket free volume for prepared experimental holo structures, the apo reference, and representative conformations selected by regular-space clustering from MCMC, backbone-restrained MD, and unrestrained MD. MCMC and restrained-MD representatives occupy the experimentally observed holo-like region, whereas unrestrained-MD representatives show substantially reduced pocket volumes.

These results associate the largest MCMC gains with experimental pocket rearrangements that contain a comparatively greater side-chain contribution. They do not identify a sharp displacement-ratio threshold beyond which MCMC ceases to improve recovery.

### Pocket Geometry Distinguishes the Sampled Receptor Ensembles

To further characterize the selected receptors, we compared pocket solvent-accessible surface area (SASA) and free volume with the experimental holo ensemble (fig. 4B; table 2). Across 925 prepared experimental structures, median pocket SASA was 975 Å^2^ and median free volume was 1076 Å^3^.

**Table 2:** Pocket solvent-accessible surface area and pocket free volume for experimental structures and regular-space representatives selected from each simulation method. Values are medians across structures.

| Technique | n structures | Median pocket SASA Å<sup>2</sup> | Δ SASA vs holo median Å<sup>2</sup> | Median pocket volume Å<sup>3</sup> | Δ volume vs holo median Å<sup>3</sup> |
|---|---|---|---|---|---|
| Experimental holo | 925 | 975.0 | n/a | 1076.4 | n/a |
| Apo | 1 | 1009.7 | +34.7 | 1094.6 | +18.3 |
| MCMC (Population) | 8 | 1041.4 | +66.4 | 1087.4 | +11.1 |
| MCMC (Regular space) | 15 | 1016.5 | +41.4 | 1087.6 | +11.3 |
| Restrained MD (Population) | 8 | 1046.0 | +71.0 | 1081.8 | +5.4 |
| Restrained MD (Regular space) | 7 | 1042.0 | +67.0 | 1095.8 | +19.4 |
| Unrestrained MD (Regular Space) | 14 | 895.0 | -80.0 | 781.2 | -295.2 |

Regular-space MCMC representatives had median SASA and free volume values of 1017 Å^2^ and 1088 Å^3^, respectively, compared with 1042 Å^2^ and 1096 Å^3^ for restrained MD. Both ensembles occupied the experimentally observed region of pocket geometry. The MCMC medians were closer to the experimental medians than those of regular-space restrained MD.

This relationship did not extend to every receptor set or descriptor. The apo structure had a SASA of 1010 Å^2^, closer to the experimental median than the MCMC median, and a free volume of 1095 Å^3^. Population-based restrained-MD representatives had a median free volume of 1082 Å^3^, also closer to the experimental median than either MCMC ensemble. Thus, improved docking recovery was not accompanied by a uniformly closer match to every aggregate pocket descriptor.

Unrestrained MD produced a distinct outcome. Its regular-space representatives had a median pocket SASA of 895 Å^2^ and a median free volume of 781 Å^3^, substantially below the experimental medians. Both unrestrained-MD ensembles showed poor docking recovery: population-based and regular-space selection recovered 0.6% and 0.4% of complexes within 2 Å, respectively, with median minimum RMSDs of 7.67 and 6.82 Å. Recovery within 3 Å remained low at 2.0% and 2.2%. Substantially reduced pocket free volume therefore coincided with the limited pose coverage of these ensembles.

## Discussion and Conclusion

Targeted side-chain MCMC sampling improved docking pose recovery from an apo starting structure on the OpenBind 2A protease benchmark. With regular-space representative selection, the MCMC ensemble recovered 74.4% of experimental ligand poses within 2 Å, compared with 51.0% for apo and 59.5% for restrained MD. These differences correspond to improvements of 45.9% and 25.0%, respectively. The lower median minimum RMSD and improvements across multiple recovery thresholds further indicate that the selected MCMC conformations accommodated experimental ligand poses more broadly than either comparator.

The relationship between recovery and experimental pocket rearrangement provides a structural interpretation of this improvement. MCMC produced its largest gains over restrained MD when side-chain displacement was relatively large compared with pocket backbone displacement. This pattern is consistent with the sampling protocol, which directly perturbs side-chain torsions while restraining the backbone near its apo geometry. Targeted torsional sampling therefore appears particularly useful for binding pockets that accommodate ligands through local side-chain rearrangements. The reduced advantage at higher backbone-to-side-chain displacement ratios suggests that greater backbone flexibility may be needed for some complexes, although incomplete side-chain sampling and docking limitations may also contribute to unrecovered poses.

Representative selection also influenced performance. Regular-space selection improved recovery for both MCMC and restrained MD, suggesting that conformations outside the most populated regions of ligand-free sampling can provide useful docking geometries. This distinction matters because the most frequently sampled receptor conformations need not accommodate the widest range of ligands. However, the regular-space MCMC ensemble contained 15 receptors, compared with seven for restrained MD, giving MCMC more opportunities to recover a reference pose under the minimum-RMSD endpoint. The population-based comparison, in which both methods used eight representatives, also favored MCMC and supports an advantage beyond this difference in ensemble size. Matched-size comparisons across several receptor counts would more clearly separate the contributions of sampling and representative selection.

Pocket geometry helps explain the poor performance of the unrestrained-MD ensembles. Their reduced free volume coincided with low pose recovery, consistent with pocket contraction limiting ligand accommodation. In contrast, MCMC and restrained-MD representatives retained geometries within the experimentally observed range. This finding supports the use of backbone restraints for preserving accessible pockets under the simulation conditions examined here, but it should not be generalized to unrestrained MD across other targets or solvent models. Moreover, aggregate pocket descriptors did not fully explain docking performance: the apo structure already had SASA and free volume near the experimental medians, yet recovered substantially fewer poses than MCMC. Detailed atomic arrangements and access to alternative conformations therefore appear important beyond overall pocket size and exposure.

The computational cost of MCMC was encouraging. The MCMC trajectory took 3.3 GPU-hours, compared with 3.7 GPU-hours for restrained MD and 19.1 GPU-hours for unrestrained MD. Ensemble docking introduces additional costs that depend on receptor count. Efficiency should therefore be assessed using the total computational expenditure required to achieve a specified recovery level.

Several features of the comparison limit attribution of the gains solely to the sampling algorithm. MCMC and MD differed in nonbonded treatment, bond constraints, hydrogen masses, and center-of-mass handling, which may influence both conformational sampling and runtime. Comparisons using matched system configurations would strengthen this attribution. The equilibrium sampling properties of the combined Monte Carlo and MD protocol were also not established; its performance here concerns the generation of useful docking receptors rather than the estimation of equilibrium conformational populations.

The docking endpoint further defines the scope of the conclusions. Selecting the minimum RMSD across all poses and receptors measures whether a near-native pose is generated, rather than whether it is ranked correctly. Self-docking consequently provides an empirical reference under the chosen protocol, not a theoretical recovery ceiling. The use of experimental ligand coordinates to define docking boxes also makes this a retrospective evaluation with a known binding region. Establishing practical screening benefits will require assessment of top-ranked pose accuracy, ligand-independent binding-site definitions, and enrichment of active compounds.

Overall, the results support targeted side-chain MCMC as a useful approach for generating apo-derived receptor ensembles, particularly when ligand accommodation involves substantial local side-chain rearrangement. Broader validation should examine additional targets, controlled backbone flexibility, matched ensemble sizes, and total sampling-plus-docking budgets. These studies will determine whether the observed improvement in pose coverage translates into more reliable prospective docking and compound prioritization.

## Acknowledgements

The authors thank Charles Xu for his advice on presentation and communication, and Erina Katoh, Sofia Passos, and Shriya Bhat, for reviewing this work. We additionally thank the Center for Excellence in Education and the Massachusetts Institute of Technology for organizing the Research Science Institute.

## Code Availability

The repository for this project is available at: https://github.com/PavSub/Improved-Molecular-Docking

## Notes

### Competing Interest Statement

The authors have declared no competing interest.

## References

(1) Amaro, R. E.; Baudry, J.; Chodera, J.; Demir, Ö.; McCammon, J. A.; Miao, Y.; Smith, J. C. Biophys. J. 2018, 114, 2271–2278.

(2) Jumper, J. et al. Nature 2021, 596, 583–589.

(3) Lewis, S. et al. Science 2025, 389, eadv9817.

(4) Abramson, J. et al. Nature 2024, 630, 493–500.

(5) Laio, A.; Gervasio, F. L. Reports on Progress in Physics 2008, 71, 126601.

(6) Sugita, Y.; Okamoto, Y. Chem. Phys. Lett. 1999, 314, 141–151.

(7) Zuckerman, D. M.; Chong, L. T. Annual Review of Biophysics 2017, 46, 43–57.

(8) Sherman, W.; Day, T.; Jacobson, M. P.; Friesner, R. A.; Farid, R. J. Med. Chem. 2006, 49, 534–553.

(9) Subramani, P.; Torsiello, J. Mol. Simul. 2026, 52, 466–476.

(10) Metropolis, N.; Rosenbluth, A. W.; Rosenbluth, M. N.; Teller, A. H.; Teller, E. J. Chem. Phys. 1953, 21, 1087–1092.

(11) Hastings, W. K. Biometrika 1970, 57, 97–109.

(12) Wang, B.; Xi, X.; Lei, X.; Zhang, X.; Cui, S.; Wang, J.; Jin, Q.; Zhao, Z. PLoS Pathog. 2013, 9, e1003231.

(13) Nelen, J. et al. The first OpenBind release: An open experimental structure–affinity dataset and benchmark for structure-based AI, bioRxiv, 2026.

(14) OpenBind Blog: OpenBind’s first release: A structure–affinity dataset for structure-based AI, https://openbind.uk/news/blog-openbinds-first-release-a-structure-affinity-dataset-for-structure-based-ai/, 2026.

(15) Eastman, P.; Pande, V. PDBFixer: An open-source package for the preparation of biomolecular structures for molecular simulations, Accessed: 2026.

(16) Eastman, P. et al. J. Phys. Chem. B 2024, 128, 109–116.

(17) Maier, J. A.; Martinez, C.; Kasavajhala, K.; Wickstrom, L.; Hauser, K. E.; Simmerling, C. J. Chem. Theory Comput. 2015, 11, 3696–3713.

(18) Nguyen, H.; Roe, D. R.; Simmerling, C. J. Chem. Theory Comput. 2013, 9, 2020–2034.

(19) Zhang, Z.; Liu, X.; Yan, K.; Tuckerman, M. E.; Liu, J. J. Phys. Chem. A 2019, 123, 6056–6079.

(20) Koes, D. R.; Baumgartner, M. P.; Camacho, C. J. J. Chem. Inf. Model. 2013, 53, 1893–1904.

(21) Wilson, E. B. J. Am. Stat. Assoc. 1927, 22, 209–212.

(22) Efron, B. Ann. Stat. 1979, 7, 1–26.

(23) Wilcoxon, F. Biom. Bull. 1945, 1, 80.

(24) Virtanen, P. et al. Nat. Methods 2020, 17, 261–272.

(25) McGibbon, R. T.; Beauchamp, K. A.; Harrigan, M. P.; Klein, C.; Swails, J. M.; Hernández, C. X.; Schwantes, C. R.; Wang, L.-P.; Lane, T. J.; Pande, V. S. Biophys. J. 2015, 109, 1528–1532.

